# THE POSSIBILITY OF AN EXTENSION OF CATARACT TREATMENT BEYOND EYE SURGERY: A NOVEL NON-INVASIVE PHOTOBLEACHING OF THE HUMAN LENS BY LED

**DOI:** 10.1101/2023.05.23.541883

**Authors:** S Desmond Smith, Bal Dhillon

## Abstract

As discussed below cataract is the world-leading cause of blindness and impaired vision with only eye surgery as a treatment. There is major requirement for an alternative which this research has provided via Genetically Modified pigs with natural cataract making live animal experiments of LED photobleaching successful monitored by fluorescence spectra. The latter were shown by EBS as spectrally identical in pigs and humans. EBS developed a miniature fluorescence spectrometer via tunable interference filters with fluorescence excited by an LED. A second LED in the blue/violet region provided a 20mW treatment beam which was shown to reverse the fluorescence signal and dramatically improve the transmission of pigs’ eyes. These results led to permission for early Human Trials. The results were judged by independent optometrists by LOCS and fluorescence in pigs and in Human Trials by LOCS and Visual Acuity. Adequate positive results have led to full length Human Trials being conducted in four European countries. Prospects of worldwide application have been therefore indicated.

## INTRODUCTION

According to WHO reports, Cataract is the world-leading cause of blindness. There are some 2.2 billion cases of visual impairment worldwide with cataract contributing around 65 million to this number. Currently the only curative treatment is cataract surgery which effectively removes the human lens and replacing it with an intraocular plastic lens (IOL). Cataract surgery is the most common surgical procedure in the UK with around 330,000 procedures being carried out each year. The formation of Cataract is closely correlated to aging and with increasing life expectancy the demand for cataract surgery is predicted to grow significantly in future. Demand for cataract surgery already exceeds surgical capacity even in highly developed nations and already exceeds availability in low to middle income countries.

The development of a non-surgical intervention which could be delivered outside of a secondary clinical care setting could provide a significant benefit to the UK and many wider healthcare systems. Edinburgh Biosciences (EBS) is currently in a world leading position in the development of a light based therapy which uses focused LED light to improve vision in individuals suffering from cataract related visual impairment. This novel treatment which allows individuals to retain their natural lens and could be delivered by trained technicians in high street opticians rather than requiring a surgeon and all of the related support.

The original inspiration for the development was a conversation between Professor Bal Dhillon (Scotland’s Senior Cataract Surgeon at the Princess Alexandra Eye Pavilion, NHS and UofE) and Smith following a successful surgical cataract lens replacement operation on Smith. Dhillon noted that current diagnostic methods for cataract were subjective and of limited accuracy. He asked if EBS could develop an objective method.

There followed a period of intense research which led to a European Union funded joint project (CATACURE). EBS developed ideas from the initial work by Kessel et al at Glostrup Hospital, Research Department, University of Copenhagen entitled “Non-Invasive bleaching of the Human Lens by Femtosecond Laser Photolysis”. This was the first publication to show that a laser light beam could photobleach a human donor lens with cataract. A small increase in transmission (5%) in a small portion of the lens was observed – indicating the possibility of significant delay in cataract development. Due to the potential of retinal damage by the laser beam, this approach was discontinued at Copenhagen. EBS recognised that in contrast with lasers the output of an LED diverges and so can be collected and focused by (say) a 50mm focal length lens to concentrate the light inside a human or porcine lens. The light then diverges to be around 60 times weaker in intensity by the time it reaches the retina. Fig. 1 shows such a ray trace.

**Fig. 1.**
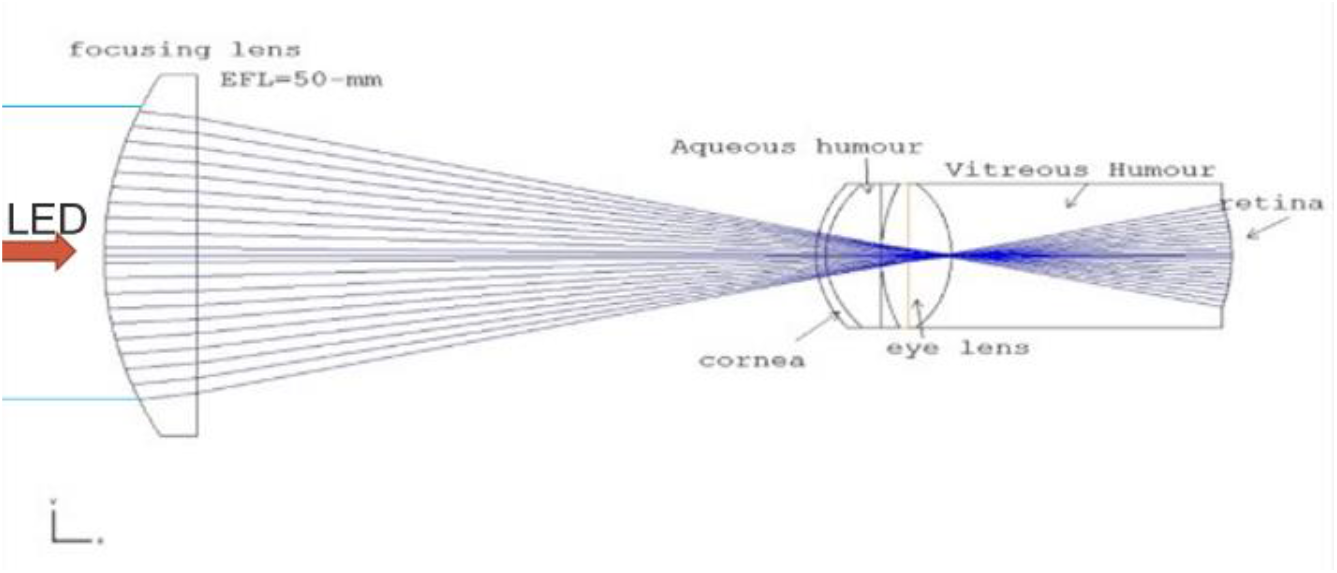
Ray Trace of spatial dispersion of treatment and excitation beams from a focus inside a human lens and expanded when incident at the retina.

It was known that UV can be a cause of cataract thus, EBS conducted early experiments on porcine lenses with cataracts induced by shortwave ultraviolet light. This enabled the timely application of the newly Japanese invented (2017) light emitting diodes (LEDs) from the semi-conductor gallium nitride. These are capable of emitting narrow band widths from less than 310nm and into the visible spectrum. (Hence a Nobel Prize for revolutionising lighting).

The absorption of light of different wavelengths by the human lens was also measured by Kessel et al in 2010 (Ref: 2). The variation of transmission with age particularly for patients likely to encounter cataract problems was determined and clearly showed lower transmission of light at shorter wavelengths Fig. 2.

**Fig. 2.**
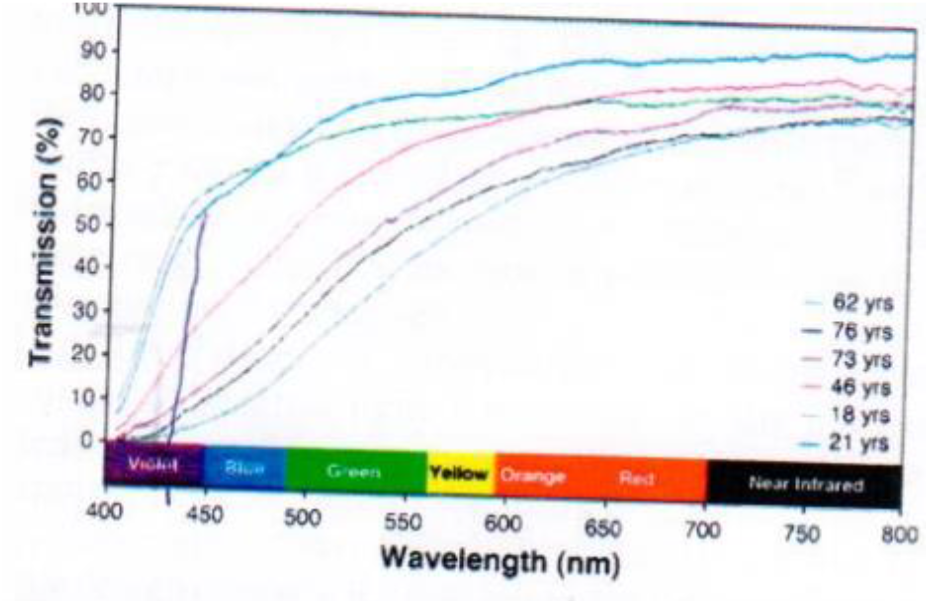
Age related transmission spectra of human donor lenses and indication of treatment wavelength.

The graph above clearly shows that at the wavelengths shown to be optimal for photobleaching (415-430nm), the transmission was less than 10% after the age of 40 years. This absorption combined with the spatial dispersion of the treatment beam implies that the energy reaching the retina is of the order of 600 times weaker than the 20mW beam energy delivered to the lens at the point of treatment. The above features provide a reasonable explanation of the safety of the treatment since no (damaging) effects have been observed in histology within the eyes in preclinical testing in pigs in numerous histology trials in Munich and Edinburgh. This is reported below in original measurements at Munich and further at the Roslin Institute, University of Edinburgh.

Responding to Dhillon’s original request to develop a diagnostic system, EBS investigated the potential of using fluorescence as a diagnostic tool. The process of fluorescence is achieved by radiating a substance to give sufficient absorption to excite the intramolecular electrons to “excited states” by using photons of energy greater than those held inside the molecules. These electrons then fall back to their normal state and emit “fluorescence light”. The result is fluorescence in wavelengths that are characteristic of different substances.

In extracted normal human lens it was shown that UV photons around 300nm resulted in a characteristic fluorescence peak at around 350nm. When the lens is affected by the formation of a cataract the 350nm band is reduced and a new peak appears at 430nm. This latter gives a potential qualitative measure of cataract. The molecular chemical change of this derivative of tryptophan is shown below (Fig. 3).

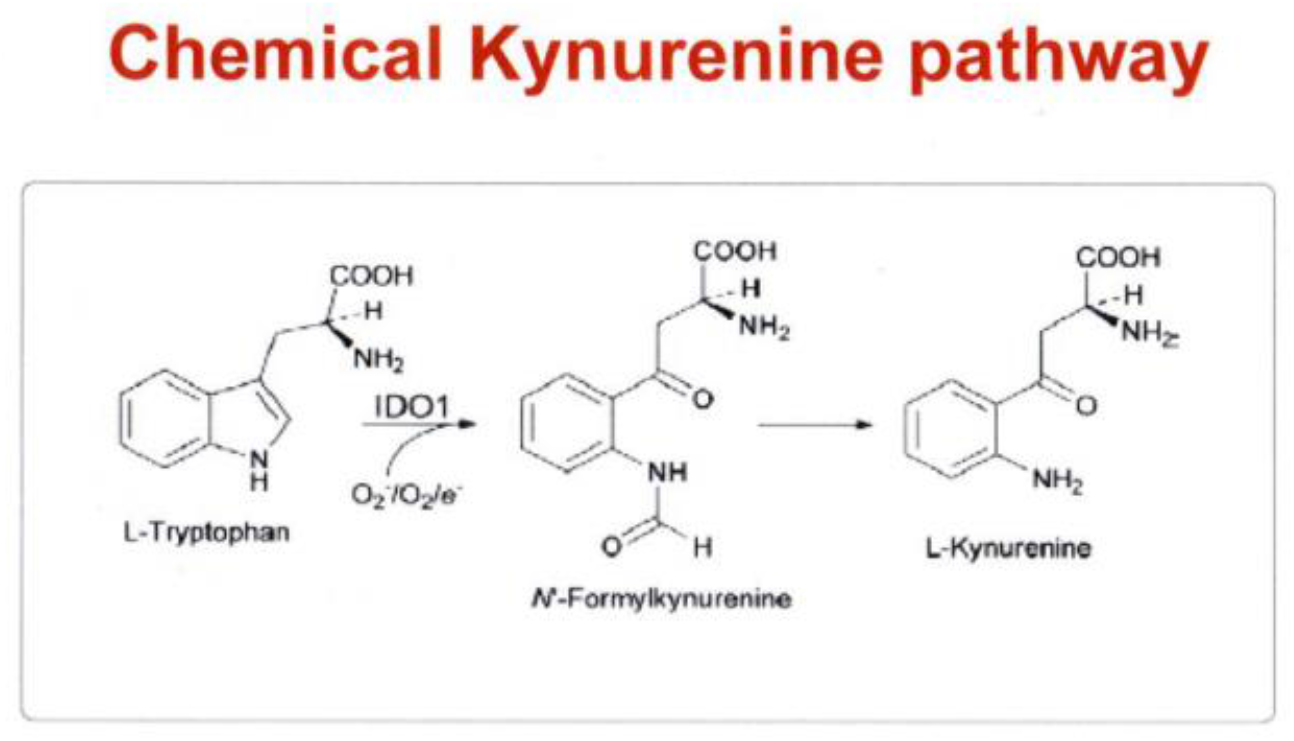

EBS had previously shown (Ref. 2: “Exploring the possibility of early cataract diagnostics based on tryptophan fluorescence”) that the fluorescence spectra of pigs’ lenses were at identical wavelengths to those of human lenses. Hence the pig’s eye makes a good experimental model. Fig. 4.

**Fig. 4.**
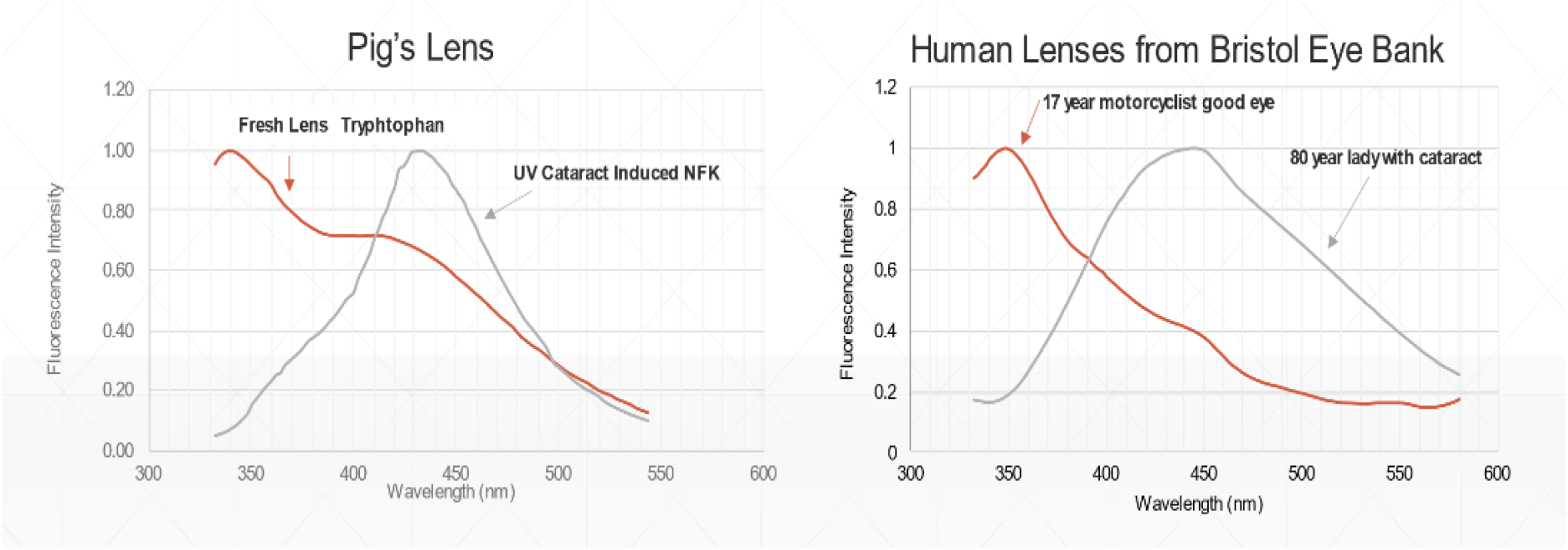
Indicating spectral similarity of pigs and human lenses.

Photographs of the pigs’ lenses were taken by the PAEP Fig. 5

**Fig. 5.**
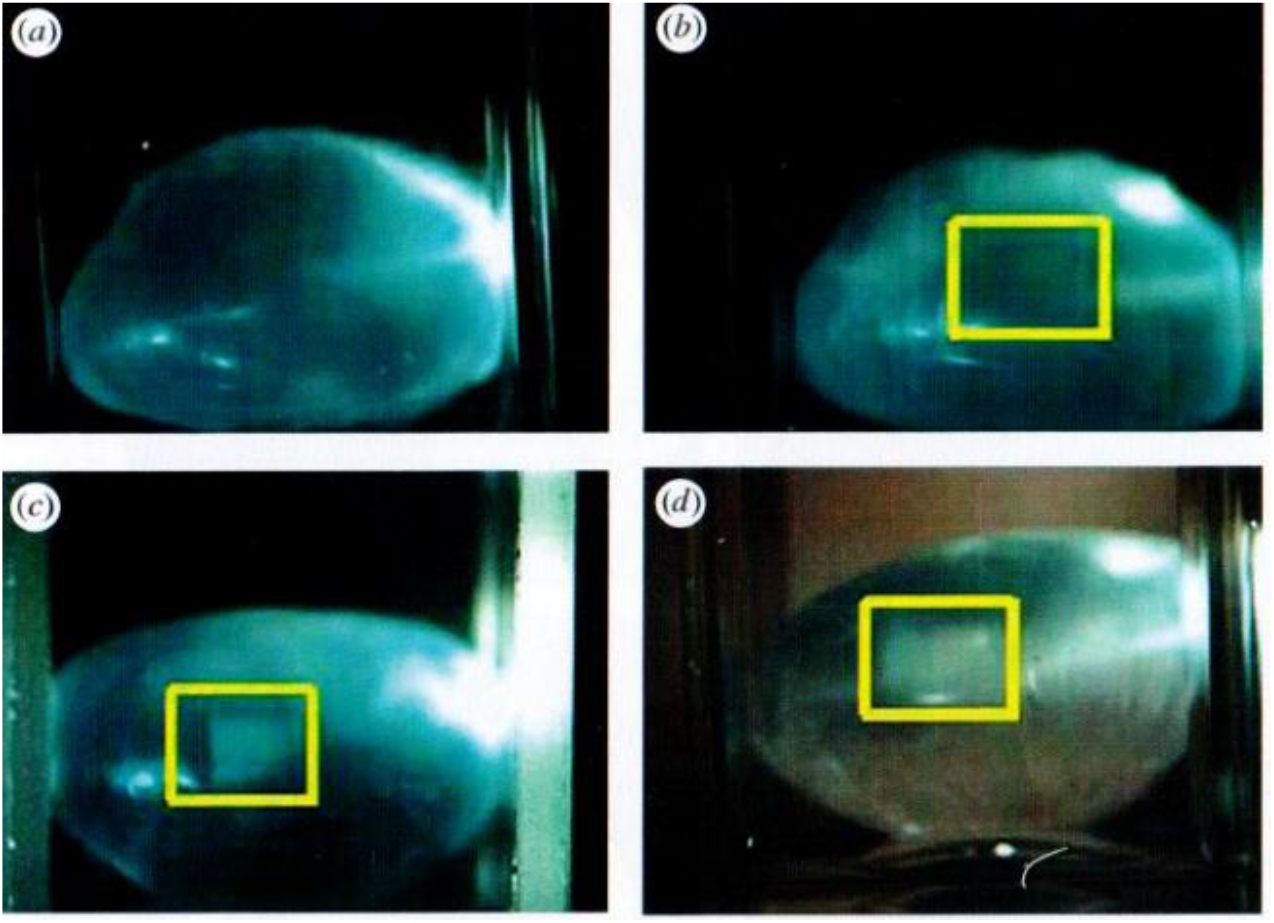
shows slit lamp images of pigs’ lenses following UV induction of cataract. Lens B remains clear although fluorescence spectra were shown in Fig. 4 changed from tryptophan at 350nm towards a band at 430nm. Scatter is only seen in lenses C and D after 4- and 12-hours irradiation. This indicates the sensitivity of fluorescence diagnosis.

### Genetically Modified pigs’ lenses

The demonstration that pig lenses were generally an acceptable model for human lenses led to extracted pigs’ lenses and whole eyes being used as initial models for research, leading eventually to treatment of live animals in association with Prof Eckhard Wolf of The Ludwig Maximillian University of Munich. The Munich team had developed a genetically modified pig strain (define) which carried genes inducing diabetes and which spontaneously develop cataracts at an early age. The previously described safety data, also carried out in pigs, suggested that these animals would be a useful modelling route to test safety and efficacy of photobleaching treatment using light at 415-440nm wavelength.

The original Glostrup laser work had deduced that the photobleaching effect was caused by non-linear optical effects creating two photon absorption around 400nm rather than the 800nm emission of the femtosecond laser. In theory the energy of this shorter wavelength is sufficient to break a number of carbon bonds such as with oxygen, hydrogen and nitrogen thus potentially affecting the molecules and causing photobleaching. Consequences were initially demonstrated by Glostrup (Ref. 3) using CW lasers in the range 400nm to 500nm with greatest treatment sensitivity observed at around 415nm to 440nm.

EBS adapted the process to create a simpler LED based optical system to focus an LED beam at 415nm into the centre of live GM pigs’ eyes for the first experiments in Munich (2018) and subsequently in a further series of studies at the veterinary Roslin Institute (2019), University of Edinburgh. The device was entitled LEDINBIO.

## MUNICH RESULTS

### The First Observation by EBS of LED Cataract Photobleaching in a Living Animal through the Cornea for both Treatment and Diagnosis

Fig. 6 shows early diagnostic spectra in blue taken with various adjustments made to the focal point of the beam on the eye.

**Fig. 6.**
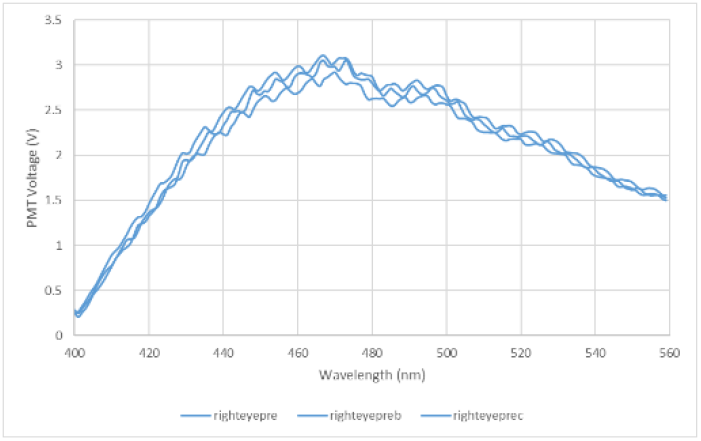

These initial spectra obtained at Munich from the LEDINBIO device described below shows regular variation due to periodic application of anaesthesia. Fig. 6.

Development of the LEDINBIO device to treat and diagnose.

### Instrument Development

The opportunity available by the creation of LEDs for the range 310nm to 430nm was combined with new developments of interference filters by Edinburgh Instruments Ltd and Delta Optical Thin Films (DOTF) in Denmark via a Eurostars grant. This created variable wavelength band pass interference filters **enabling** by combination of LED excitation to create a miniaturised fluorescence spectrometer and treatment device entitled LEDINBIO. This device in laboratory form is shown in Fig. 7 and was combined with a Keeler slit lamp microscope in Fig 7. This, Point of Care capability was able to be confirmed with the size indicated in the diagram.

**Fig. 7.**
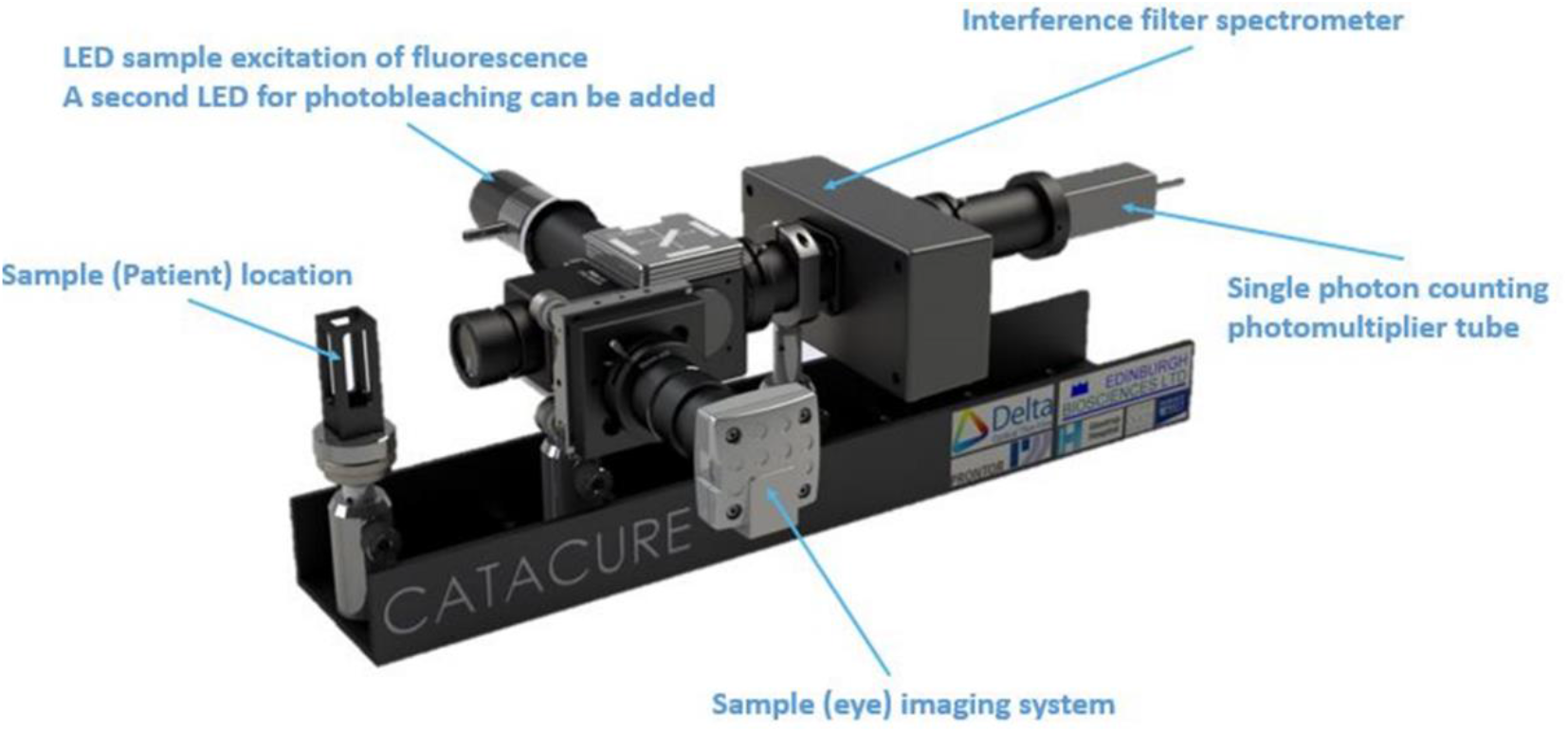
Lab version of LEDINBIO: Important dimensions: length 35cm breadth 12cm/15cm (the scanned filter) height 10cm.

A small USB connected computer-controlled circuit capable of driving the LED at variable power and decoding the received signal using phase sensitive detection to reduce background interference was designed and constructed.

DOTF found that this application of variable interference filters could also be combined with semi-conductor diode array detectors. Consequently there are two methods to obtain the miniaturised spectra. Firstly, to translate the variable filter across the entire fluorescence beam received by the LEDINBIO equipment (which had high light grasp and high sensitivity with photomultiplier detection). Secondly, the combination of the variable filters with the semi-conductor arrays can also give a spectrum but starting signals will be reduced by the number of elements in the array and such diodes are not as sensitive as photomultipliers hence the overall device in that form is less sensitive.

LEDINBIO is a combination of **LEDs with Tunable Interference Filters** and focusing optics and photomultiplier detection providing both treatment and diagnosis in a small instrument.

### Experimental Arrangements

The anaesthetised pigs were simply laid on a bench at around one meter height. Simple optics were arranged between the treatment and fluorescence excitation sources – LEDINBIO. The LEDs were mounted on a microphone stand adjustable in height and position to illuminate a pig’s eye via a lens and or fibre. These rays included together with a beam splitter were manually adjusted and served also to collect the fluorescence return from the pig’s eyes.

The first observation Fig. 8 scanned from 400 to 560 by motion of the variable interference filter of width 10nm (Fig. 7). The oscillations on Fig. 3, 8 and 9 were due to the periodic injection of anaesthetic on the pig and could reasonably be averaged out.

**Fig. 8.**
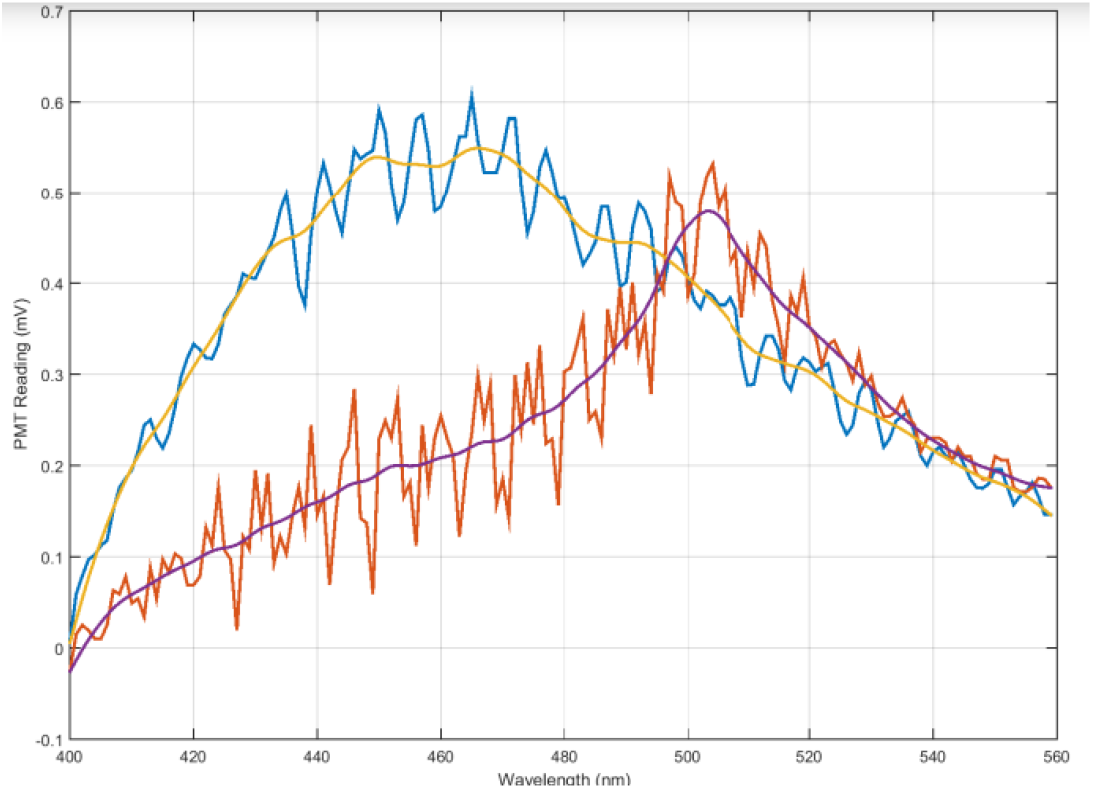
The blue curve was the GM pig before treatment but with the control pigs’ background fluorescence subtracted. The red curve shows the diminution as a result of treatment and also a change in spectral form. This is an important result increasing the sensitivity of detection. Fig.9 with the oscillation averaged.

**Fig. 9.**
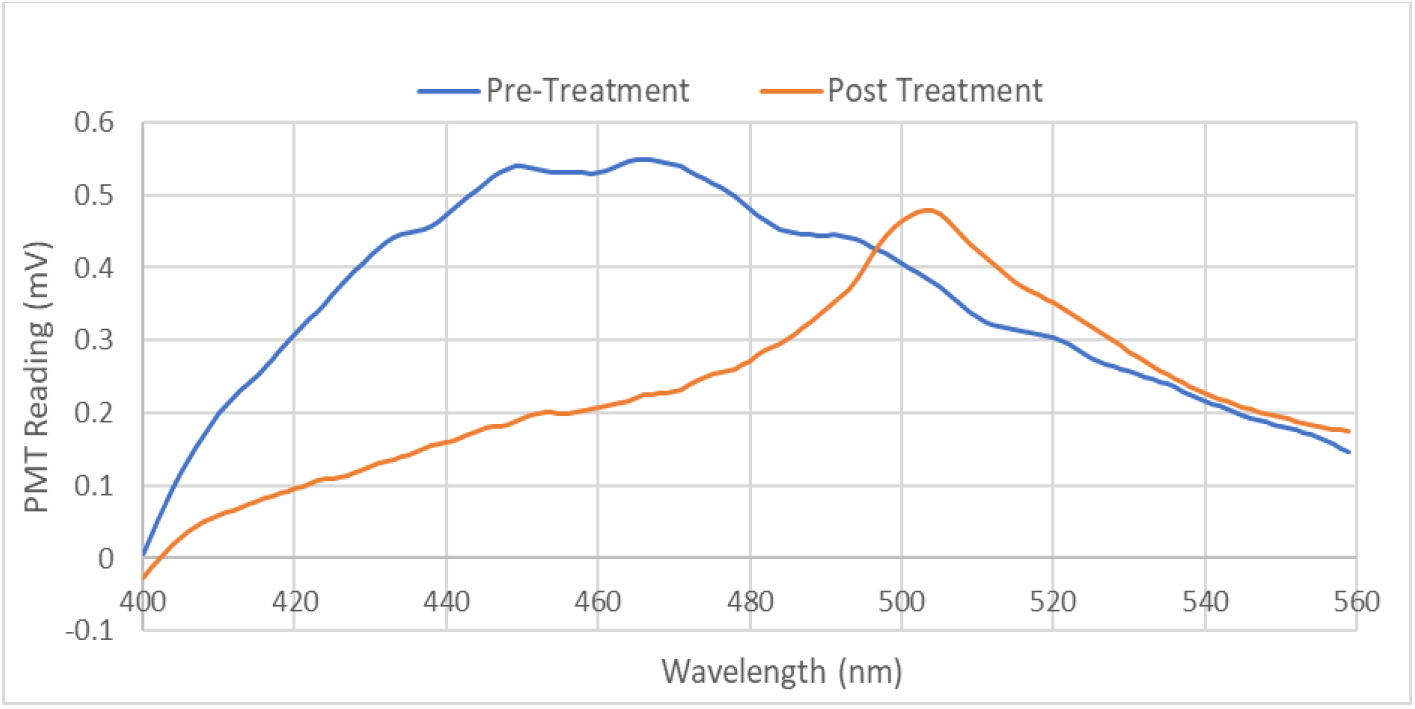
INSC94Y transgenic pigs: Pigs 6025, 6026, 6027 Commercial pigs: Pigs 6023, 6029.

### First Photobleaching Result in Munich

These initial studies on living animals (therefore including the cornea) were made with a total of 5 eyes in 3 GM animals and 4 eyes in 2 commercial animals. In 4 eyes (4 animals), a treatment power of 24mW was studied, delivering a total energy of 172 J. Fig. 3 above shows how the regular variations coincided with the periodic application by the veterinarians of the pigs’ anaesthetic affected the fluorescence output. We can reasonably average it as shown.

Following such analysis the curves became as in Fig. 9.

These Munich results with background subtracted before and after treatment show a **Decisive Reduction following photobleaching** and change of spectral shape. The spectral chemical properties of various kynurenines were reported during CATACURE by Ref. 3 Gakamsky et al “Scientific Reports” 2017.

The background is due to various kynurenines i.e. fluorescence even if complete photolysis does not go to zero. The result is therefore a clear change after treatment in both intensity and spectral change (the latter could be an important effect independently from absolute fluorescence levels) – an important future procedure suggesting the need for availability of controls in Human Trials and determination of the corresponding “zero level” in normal human eyes will be required. Since the normal eye fluorescence is significantly smaller than say a grade two cataract absolute accuracy will not be necessary to obtain a reasonable quantitative level of cataract by fluorescence.

Histology of these eyes, was performed by a veterinary pathologist at Munich and found no evidence of damage, such as retinal scars or typical lesions and demonstrated the proof of concept for treating cataract lenses with the LEDINBIO device. Therefore, it was considered that the input energies and exposure times were safe and it was acceptable to proceed towards Human Trials.

### Detail of Cataract Formation

The visual impairment of cataract is due to both increased absorption and scatter in the human lens. The origins have been illustrated by Fig. 10.

**Fig. 10.**
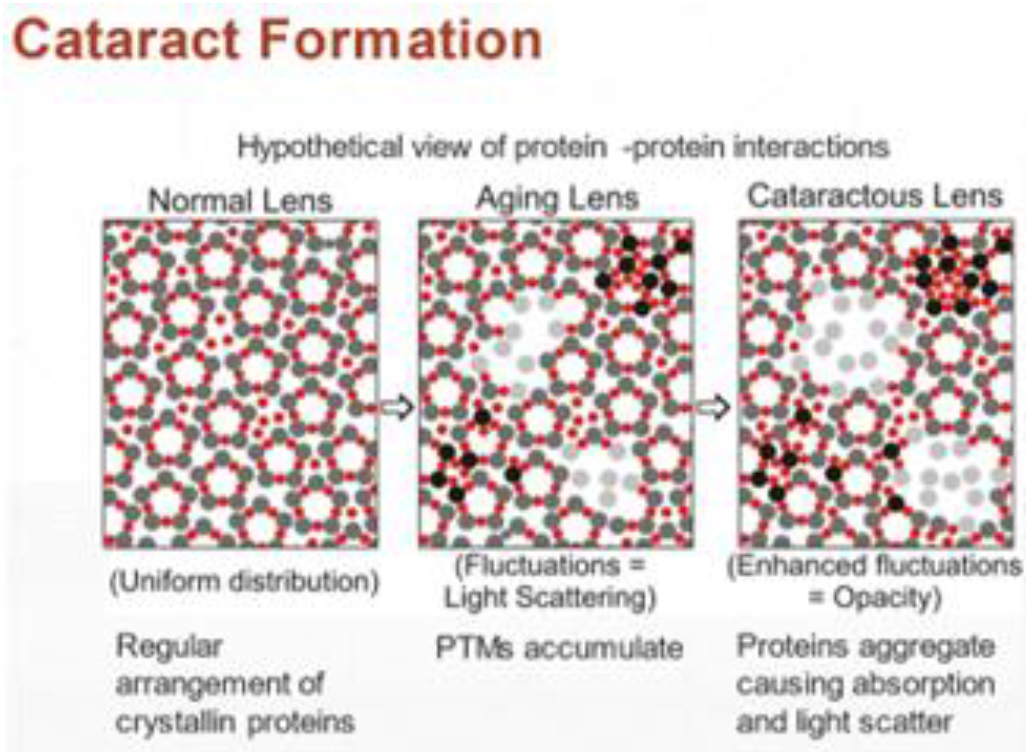
Ref. 7 Role of short-range protein interactions in lens opacifications. Mol Vis. 2006 Aug 10;12:879-84. PMID: 16917488.

**Fig. 11.**
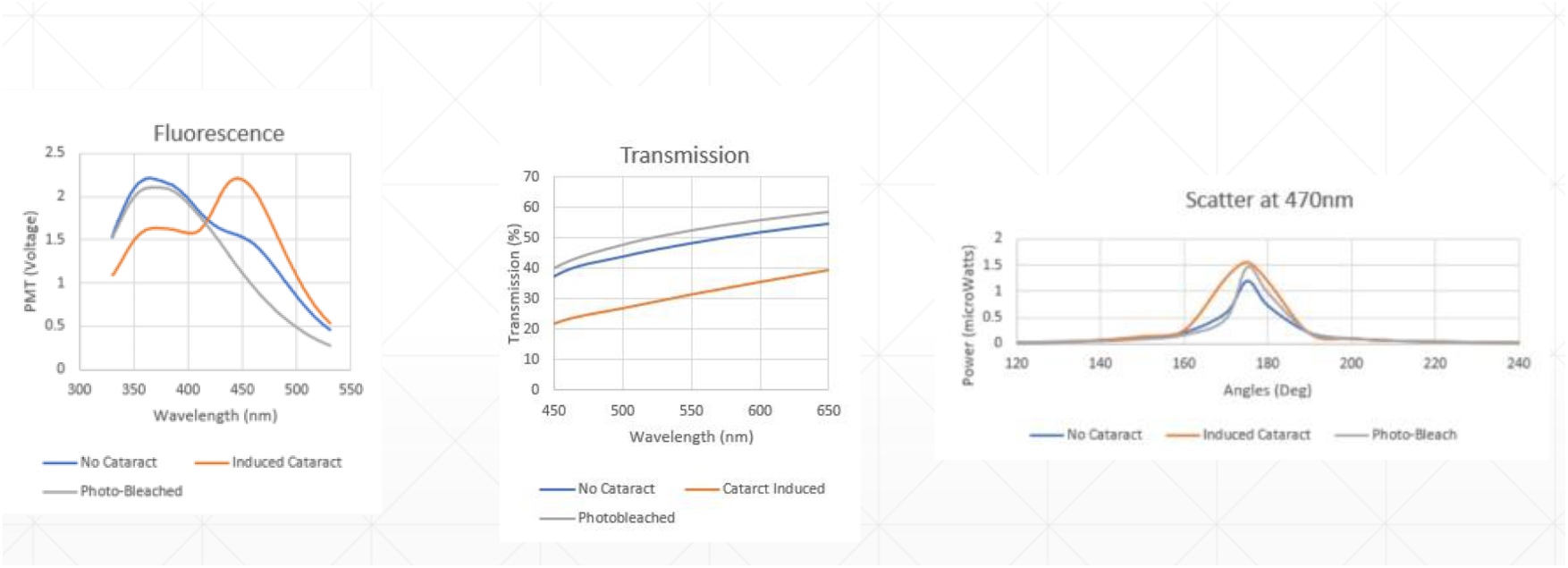
The fluorescence spectra of extracted pigs’ lenses were recorded by the LEDINBIO Interference Filter Spectrometer and followed the typical format as shown above.

### Correlation between Fluorescence, Transmission and Scatter

The results (in Fig. 12 below) were achieved with extracted pigs’ lenses that showed a correlation between the fluorescence changes and changes in transmission and scatter. Such results have been repeated on a number of samples.

**Fig. 12.**
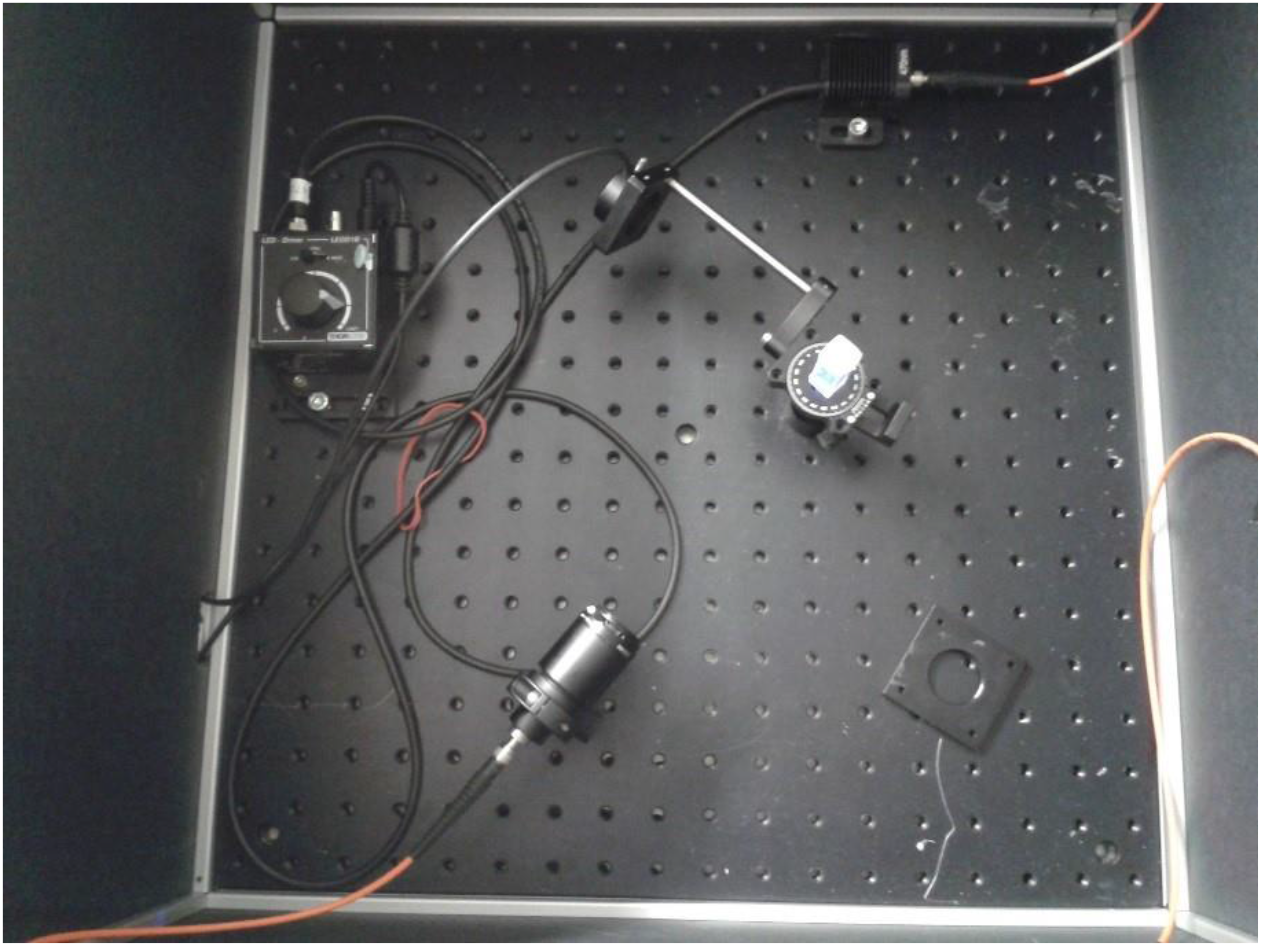

### Correlation

Creation of cataract was by UV exposure for 2 hours at 310nm and this caused the NFK spectrum. Photobleaching for 3 hours at 420nm (11mW, focussed on a 1–2mm diameter spot) caused the spectra to revert to something close to the original.

Transmission had earlier been measured by Line Kessel et al Fig. 2, Ref: 3 using a Xenon lamp with a collimated white light beam 3mm diameter. After the sample lens was an integrating sphere collecting the transmitted beam to feed into a USB spectrometer. The results showed a similar pattern to the fluorescence spectra with the **photobleaching increasing the NFK transmission** returning to something close to the initial figure.

The transmission measured from the photobleached lens was slightly greater than that recorded for the fresh lens, this may be due to some surface smoothing under photobleaching.

The Scatter results were recorded using a rig constructed in the lab, photograph shown in Fig. 12.

### Angle Scattering Apparatus

The 470nm, 6mm diameter source beam illuminates the front face of the test lens and the detector is rotated around at 120mm radius, detecting the power of the scattered radiation. The detector only moves in the horizontal plane.

It should be noted that both the transmission and scatter experiments sample an area larger than that directly affected by the UV-irradiation.

The scatter results show that most of the light passes straight through the lens, as expected. The original width of the peak is a measure of the forward scatter, and the Full Width Half Maximum of the peak recorded for the induced cataract treatment is approximately double the width of the peaks. This is significant because this is the scatter ‘seen’ by the patient. The blurry effect described by cataract sufferers is the result of forward scatter of light within the eye.

## Conclusion of Scatter

These results prove that the photobleaching does improve both the transmission through the lens and reduce the scatter that adversely affects vision. This is important as it is the most severe cataracts are dominated ultimately by scatter and we show that this is removable by the LED Treatment. A possible mechanism is noted above.

### Experiments at the Roslin Institute, University of Edinburgh

These experiments were repeated using 365nm LED excitation for fluorescence to provide more numerous results with better incident optics. They operate through the cornea with a satisfactory transmission wavelength.

These experiments were on 18 complete GM pigs’ eyes to overcome any difficulties from other ocular components. Extensive histology indicated no damage to further eye components. These results were possible through the EU CATACURE programme 2015-2018.

The final state of cataract depicts substantial aggregation of the proteins which are responsible for the scatter.

The Roslin results confirm the Munich results with some 18 GM pigs which were imported from Munich to the veterinary facilities of the Roslin Institute. An improved optical system discussed below was used to determine the optical fluorescence spectra of which a typical result is shown below. Fig.13

In all the 18 cases at Roslin independent assessment before and after treatment were **made by professional optometrists** using the standard LOCS method. Although Fig. 12 above typically does not show the fluorescence going to zero after treatment. Further assessment of visual acuity on the LogMAR scale was of course not available from pigs. Nevertheless, on pig no. 2 (not the spectrum of Fig. 13) of the series **Optometrist, Pam McClean epo ted “Pig 2 was the most ema kable from my perspective. There was a definite grade 2 present in the right eye. Post t eatment the ight lens was completely clea “**.

**Fig. 13.**
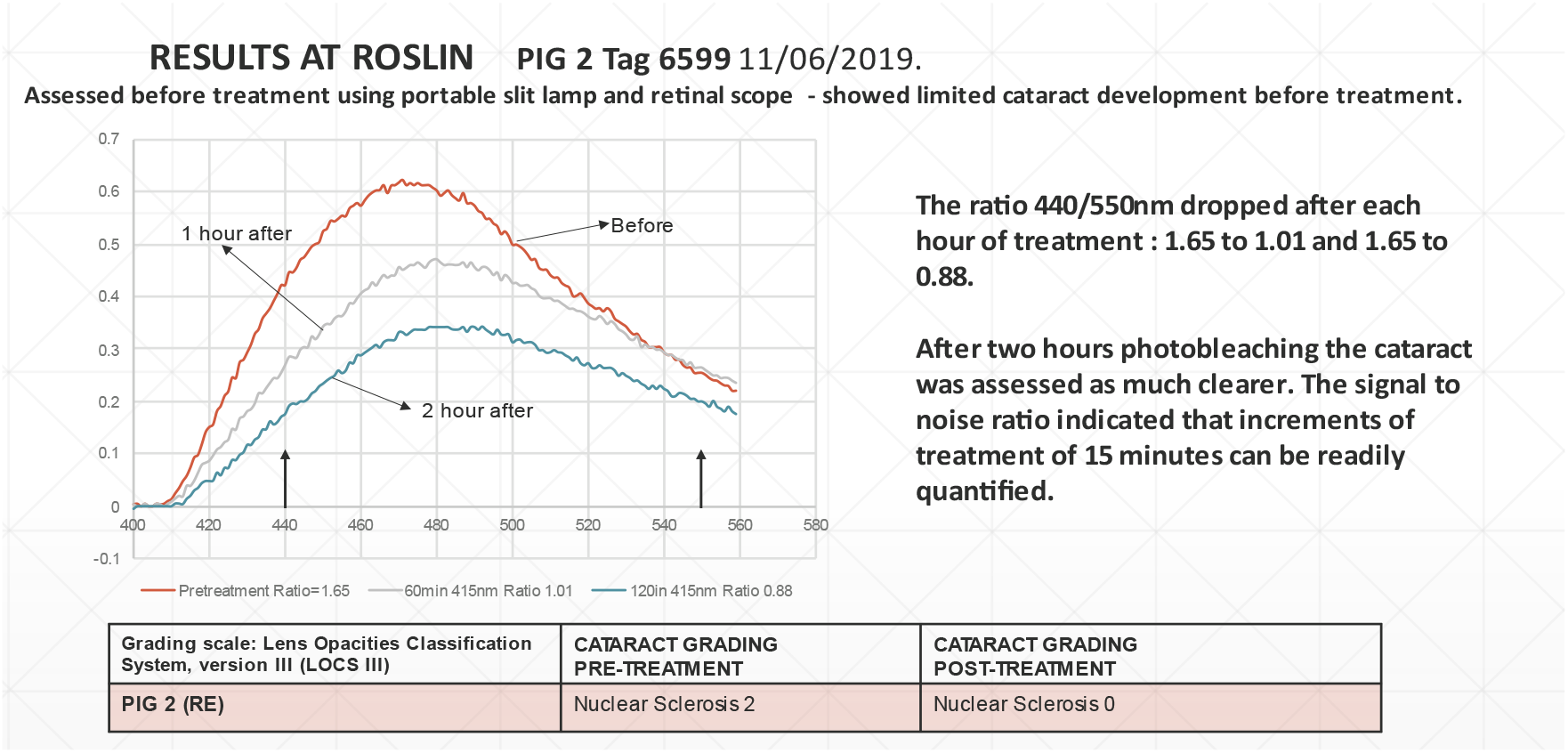
Spectra from Roslin (showing non-zero fluorescence following treatment).

This suggests that there was a complete cataract clearance even though no background (due to further kynurenine molecules) was available for subtraction.

The combination of the Munich and Roslin results followed by the EBS creation of the LEDINBIO Diagnosis and Treatment device enabled first Human Trials in Latvia to be considered by Onorach the CRO from Dundee.

### Possible Mechanisms of Scatter

When cataract worsens the absorption effect leads to scatter and the lens appears white. As stated in the pictures above, this is due to the formation of protein-protein aggregation. The white colour observed implies that the scattering aggregates are, (by Gustave Mie electromagnetic theory), roughly the same size as the wavelength in the visible, i.e. around 500nm. **These aggregates will only scatter if their refractive index differs from the background**. Following the change of transmission (and projected into the UV) the Kramer’s Kronig relationship implies through the integral of the UV absorption controlling the refractive index that our treatment is probably reducing the aggregate refractive index and hence the scattering itself. This remains to be quantified but is almost certainly a part of the process. The scattering results are shown in the right-hand Fig.12 which shows a non-scattering optical system angle expanded by a factor of more than 2 with cataract and reducing almost entirely to the original following LED treatment. This is a very important result that not only the molecular changes controlling absorption but the ultimate scattering which impairs visual acuity can be effectively treated.

### Instrument Development for First Human Trials

To enable the first Human Trial EBS made a **combination of LEDINBIO with the Keeler slit lamp microscope**. Fig. 14

**Fig. 14.**
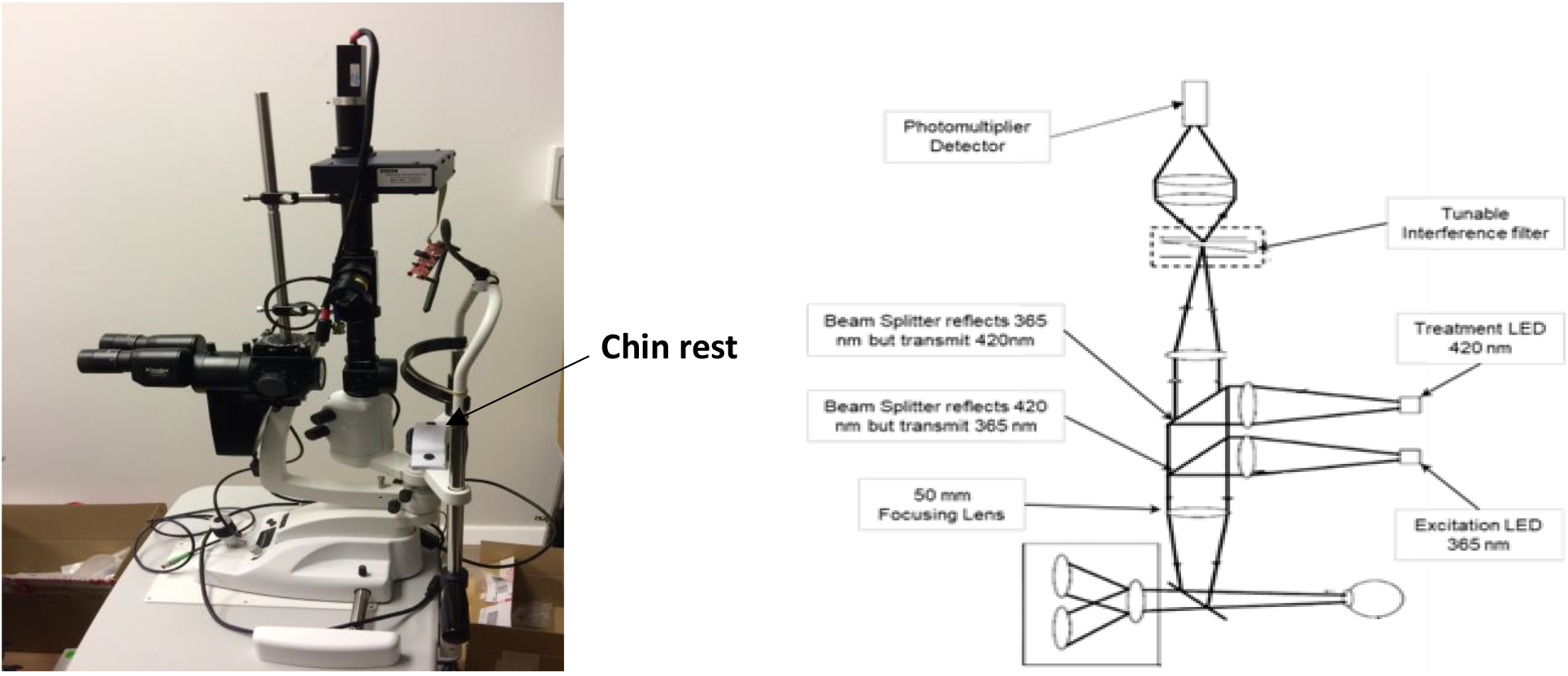

**Fig. 15.**
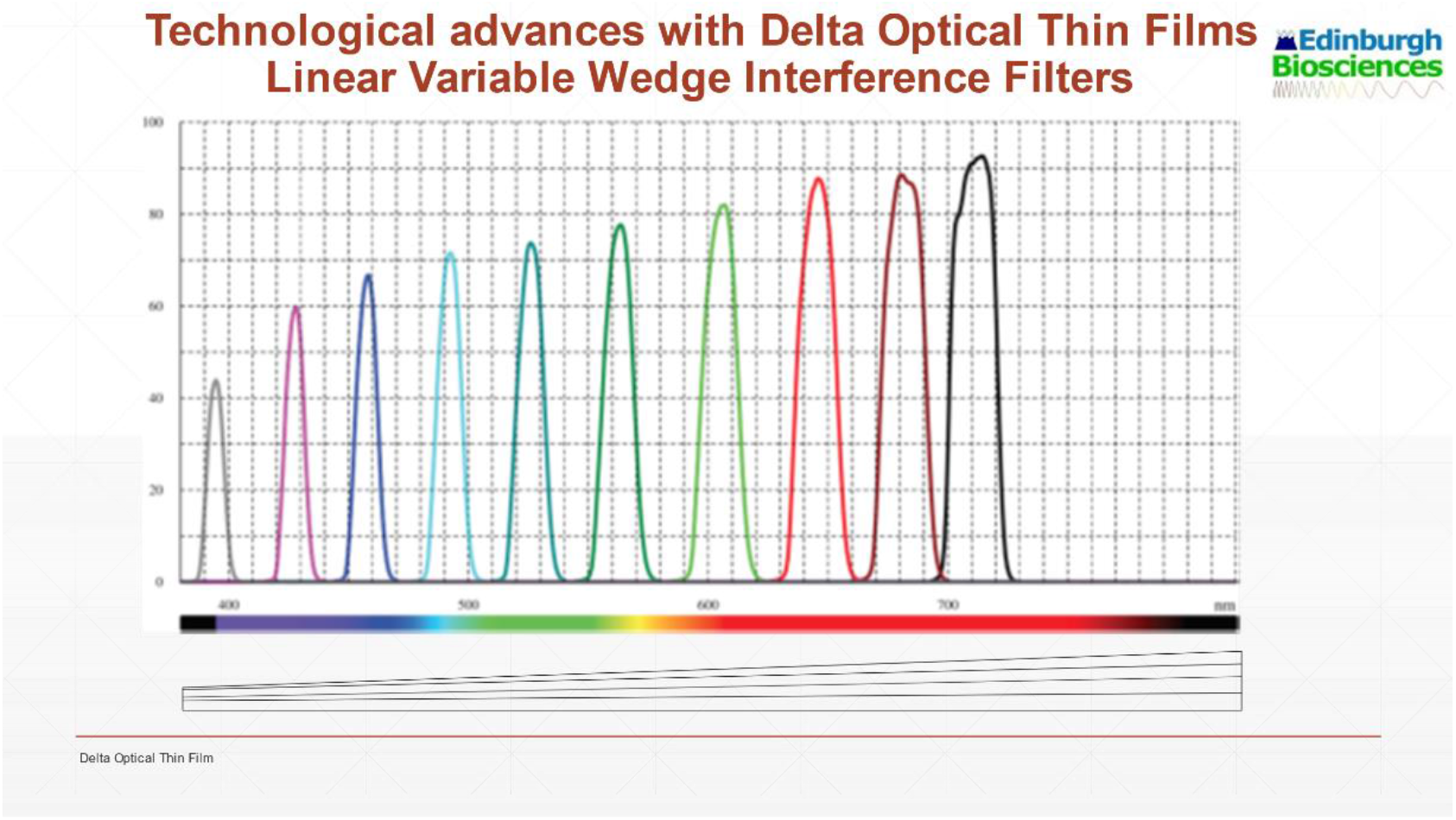
Transmission Spectra as moved along the wedge.

This non-invasive photobleaching enabled a First Human Trial in Latvia of 11 patients with cataract grades 1, 2 or 3. **In results all patients showed evidence of improvement in the LOCS assessments**. With humans as the patients, it was also **possible to assess visual acuity** which was not available from the pig

Diagnosis and Treatment of humans proved possible by fluorescence using an LED at 365nm for excitation of the band at 430nm. Such fluorescence bands are broadened to the extent of about 40nm due to the thermal vibrations of the complicated molecules. The chemical kynurenine pathway was shown in Fig. 4.

Fig. 4 identifies the natural healthy eye producing a band at 350nm. This transforms to the band of N-Formyl Kynurenine (NFK) which is a degradation product in the conversion of tryptophan to kynurenine.

The use of fluorescence spectra for diagnosis requires the EBS creation of the **miniaturised fluorescence spectrometer** capable of **Point of Care (POC) scale of use**. EBS has created such an instrument following joint work between Edinburgh Instruments Ltd and Delta Optical Thin Films in Denmark with a Eurostars Grant of € K.

This need to create a small spectral instrument with high light grasp follows from S D Smith’s original paper, Ref. “Design of Multilayer filters by considering two effective interfaces” 1957. **Such filters have high light grasp and can be made wavelength tunable by depositing all the multilayers in the form of wedges**. Thus, a derivative of Smith’s double half wave narrow band filter was prepared by Delta Optical Thin Films covering wavelengths from 300-600nm with a band width approximately 10nm. **By focusing the collected fluorescence return from the human eye to the order of 1 or 2mm and moving the variable filter across the beam the band width is retained and a complete spectrum obtained**. Such filters have a high light grasp and can be used with the N-Formyl Kynurenine (NFK) cataract detection band. This is possible without loss of effective spectral resolution since such NFK bands are automatically broadened by molecular motion of the complicated molecules to a width around 40nm or 50nm. Thus, a highly sensitive miniature fluorescence spectrometer was created and scans with a piezo electric stick and shift motion as illustrated in Fig. 16.

**Fig. 16.**
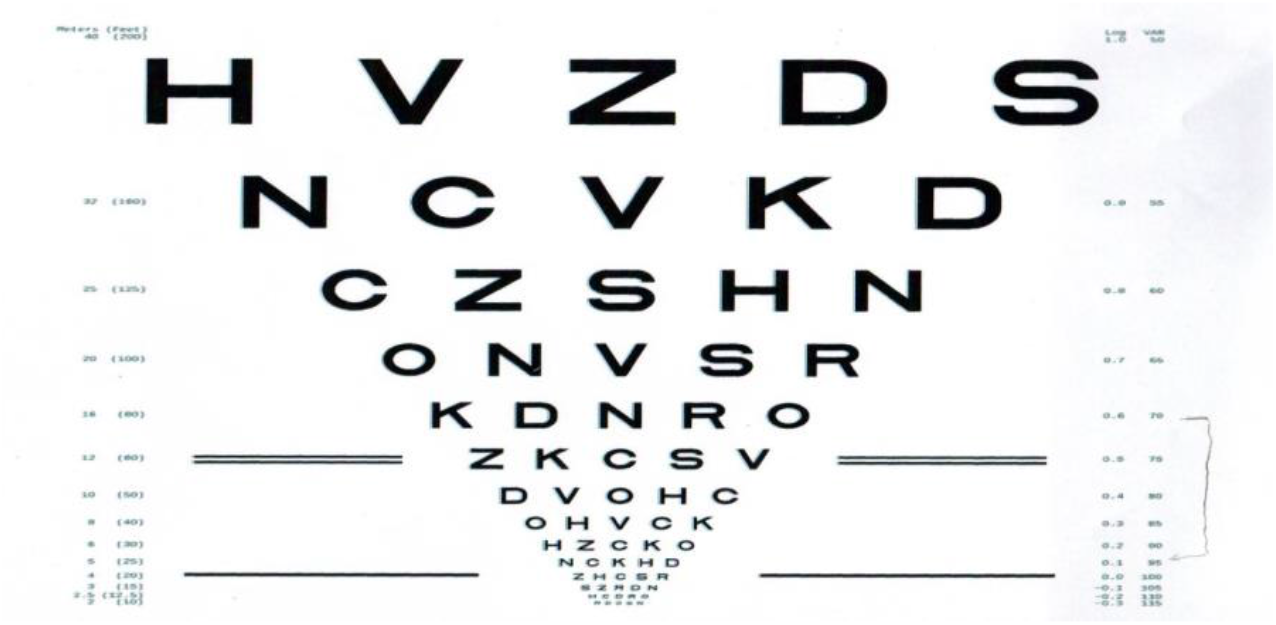

A variant is to use the variable filter in combination with a detector array which can give the complete spectrum instantaneously but with considerable reduced sensitivity by spreading the total fluorescence spectrum individually on to the 50 linear elements of the array.

This also illustrates the use of phase sensitive detection to eliminate any background noise as required by the small fluorescence signal. A diagram of the optical and mechanical system is given in Fig. 16. The above instrument was used to detect the early signs of cataract with a European Union Eurostars grant to create practical tunable interference filters.

The variable filter output is electronically treated to deliver signals from successive elements creating the spectrum.

The LEDs themselves have a natural band width of emission with about 15 to 20nm half width and the exact wavelength for treatment is at least as wide as this. EBS combined the Miniature Fluorescence Spectrometer with the treatment LEDs with directive and focusing optics as LEDINBIO in Fig 6 above. This was combined with a Keeler microscope containing directional and angular adjustments to project and receive from a live human eye. Thus, suitable equipment was created for a First Human Trial.

## RESULTS OF HUMAN TRIALS

### A summary of the First Trials in Latvia & Lithuania

In order to proceed to first human trials EBS created the first and necessary Investigators Brochure and together with the CRO, Onorach, an appropriate Protocol. As a first Proof of Concept there were 11 patients (a complete report has been made by Dr Crispin Bennett). One notable result was a patient who had lost his driving licence due to visual impairment and following treatment under the Protocol was able to regain it and recommence driving. Compared with the pigs’ results humans were able, of course, to provide visual acuity. One such result at the beginning of further trial recorded a starting LogMAR of 0.6 to after treatment 0.1 (Fig. 17)

The success of the first trials is indicated by results by independent optometrists of LOCS and LogMAR visual acuity results.

### Further Trials in the Baltic States

A second trial has currently commenced in Lithuania. This shows a significant result photobleaching in eight 15 minute visits a grade 3 cataract with Visual Acuity initially at LogMAR 0.6 to conclusion at visit 8 when LogMAR was 0.1.

## Conclusion

The LED treatment can be decisive with powers of the order of 20mW and 2 hours of total treatment. Due to the early work of Glostrup on transmission as a function of age where cataract is obtained a series of animal experiments with pigs showed the transmission at the effective treatment wavelength was less than 10% across a range of age and since the beam was focused in principle on the back surface of the lens capsule then reduced by a factor of 600 by the time it reaches the retina. Thus, it is established that the treatment does not damage other parts of the eye. Thus, the conclusion is that the LED/fluorescence route to treatment and diagnosis is a practical proposition which could be operated by many opticians worldwide and would alleviate the surgery problem and consequent waiting lists.

### Patents

EBS has been granted patents in USA, Europe and Australia with ongoing application elsewhere.

## Acknowledgements

We thank Professor Bal Dhillon and the Princess Alexandra Eye Pavilion, Professor Eckhard Wolf, Delta Optical Thin Films, Heriot Watt and Edinburgh Universities for varied assistance and contribution. Grants have included partial contributions from Eurostars, NHS (NIHR), MRC, European Union CATACURE, Innovate UK and substantial remaining support from EBS itself.

